# AKT-*like* kinase promotes cell survival during nutritional stress in trypanosomatids

**DOI:** 10.1101/2020.07.25.218644

**Authors:** Andrés Felipe Díez Mejía, María Magdalena Pedroza, Lina M. Orrego, Sergio Andrés Pulido Muñoz, María Clara Echeverry, Mauricio Rojas, Maurilio José Soares, José María Pérez-Victoria, Carlos Enrique Muskus, Sara María Robledo, Marcel Marín-Villa, Rubén E. Varela-Miranda

**Author notes:** These authors contributed equally to this work (Marcel Marín-Villa, Rubén E. Varela-Miranda).

## Abstract

Tritryps are protozoan parasites that belong to the Trypanosomatidae family, which encompasses the etiologic agents of leishmaniasis, African and American trypanosomiasis. These parasites undergo different stress conditions across their life cycle, such as nutritional stress, which needs to be deadened in order to guarantee the survival of the parasite inside its vector and mammal hosts. Here we show that the lack of the serine threonine kinase PKB / AKT-*like* function, either by allosteric inhibition of its Plekstrin domain (PH) in *T. cruzi*, the reduction of the gene transcripts in *T. brucei* by RNAi assays, or by AKT-like gene knockout in *L. major*, reduce the tolerance to nutritional stress of parasites that triggers apoptosis-like events, including DNA fragmentation, mitochondrial damage and loss of plasma membrane integrity. Additionally, we observed that double knockout of Akt-*like* in *L. major* impairs its infective capacity. This work confirms some of the previously described functions regarding parasite survival for AKT-like kinases in the *Leishmania* genus. The present work also provides strong evidence of the probable function of Akt-*like* in *T. cruzi a*nd *T. brucei* survival and infectivity.

**Author summary:** Endemic countries for neglected tropical diseases are called to play a paramount role in the discovery of new drug candidates through the application of new drug development strategies. Rational drug design method have proven to be compatible with the development of new drug for orphan and neglected diseases since it substantially reduces the costs of discovery and development, a desirable condition for public funded initiatives. Previously we have identified a new parasite protein kinase (AKT-*like*) as promising new target candidate by means of computational tools and probed its biological role in trypanosomatids. Here we show that inhibition of the AKT-*like* kinase in trypanosomatids by different approaches (chemical inhibition, interference RNA and gene knockout) decreases the fitness and survival of the parasites *in vitro*, interfering with the capacity of the parasites to react and survive stress conditions similar to those experienced by the cell in the natural life cycle. Additionally our results strongly supports the potential of a new family of compounds previously described by bioinformatics means as potential trypanocidal agents. Altogether we show that the specific inhibition of the AKT-like is a promising strategy for the further development of anti-trypanosome drugs.

## Introduction

The World Health Organization (WHO) recognizes the African trypanosomiasis, the American trypanosomiasis and the leishmaniasis as orphan diseases caused by the parasites of the Trypanosomatidae family: *Trypanosoma brucei, Trypanosoma cruzi* and *Leishmania* spp., respectively, which are collectively known as Tritryps. Tritryps members have complex life cycles that involve the development of differential stages in mammal hosts and vector insects and the coordination of their cell cycles for the development of extra and intra cellular forms with highly replicative to non-replicative infectious behaviors [1–5].

Up to date, there is not an effective vaccine against those pathogens and the available treatments for over 50 years are based on chemotherapeutic strategies using pharmaceutical entities with rather little knowledge about their mechanisms of action. This uncertainty has led to the lack of understanding about the resistant mechanisms raising in several parasite populations around the world and, more serious, the toxic effects those drugs exerts in treated patients [6]. In that sense, the search for new therapeutic targets is a promising strategy for rational drug design for orphan tropical diseases where the knowledge of the mechanisms of action of new chemical entities is of utmost importance in order to reduce the risk of side effects and increase the chance of therapeutic success. Consequently, the kinases represent a vast source of promising targets for the development of new medicines due to their involvement in the maintenance of the cell homeostasis. They are often reported as essential and preserved throughout the eukaryotic evolution and several approaches of chemical kinase inhibition have proven to lead to cell death processes [7–10].

In 2005, the *T. cruzi*, *T. brucei* and *L. major* kinomes were published, showing a kinase network responsible for the signal transduction involved in the regulations of the cell cycle and the responses that these parasites must stablish against the adverse environments they must face in their hosts and vectors [11,12]. The Tritryps genomes have almost 30l.jl,% of the kinases shared with the human genome, occupying approximately 2% of the genetic inventory in the parasite. The Tritryps kinases have been classified in two groups named typical (eKPs) and atypical kinases (aKPs), as it is in other eukaryots [11].

Some serine/threonine-like kinases and phosphatases such as the phosphoinositol kinases (PIKs) are described as regulators of the cell survival by means of the inhibition of the apoptosis, the activation of the energetic metabolism, the immediate response to stress stimuli and the control of the cell cycle [8,13]. The PIKs kinome of *T. cruzi, T. brucei, L. major, L. braziliensis* and *L. infantum,* published in 2009, concluded that these kinases might behave as promising therapeutic targets. The PIKs were classified in 5 function models as previously proposed for *Schistosoma mansoni* according to their sequence homology, the domain similarity, the substrate specificity, and their particular regulations [14–18].

Model 1 PIKs are involved with the cellular transport and recycling; model 2 PIKs like RAC serine/threonine kinase (AKT) are related with cell survival and downstream kinase activation; models 3 and 4 PIKs are associated with the activation of the MAPK and the regulation of the cell cycle; and finally, the model 5 which are mediators of interactions between proteins and PIKs related to the DNA breakdown-dependent activation and DNA repair pathways [16,17].

AKT connects signaling routes of the PIKs/AKT/mTORc pathway and has three conserved structural domains: An N-terminal pleckstrin homology (PH) domain which binds to a PI3P, a catalytic kinase domain that is fundamental for the phosphorylation of its substrates and the regulatory C-terminal domain, which regulates the activity of the enzyme [19–23]. In humans, the AKT is located in the cytoplasm and is recruited to the cell membrane through its PIP3-PH binding domain after the formation of PIP3 in response to stress. This event promotes conformational changes of the protein leaving it available for the further phospho-activation by kinases such as PDKs or the mTORc2 complex. Once activated, it will perform a series of secondary kinase phosphorylations that regulate processes such as the inhibition of apoptosis, growth, protein synthesis, proliferation and autophagy [16,17,20–23].

We recently reported the ORFs encoding putative AKT (AKT-*like*) in the genome of trypanosomatid parasites. These ORF sequences present the 3 characteristic structural domains [24]. We further showed that in parasites of the genus *Leishmania*, this kinase (GenBank/EMBL accession no. XM_001684621) is phospho-activated under conditions of thermal and nutritional stress, suggesting a role in cell survival processes [25]. Additionally, we published evidence supporting that the chemical inhibition of the function of this protein in parasites under stress conditions induces apoptosis-like events and impairs its infective capacity [24,25]. Here we present further evidence of the importance of the AKT-*like* protein inside trypanosomatids and evaluate, for the first time, the role and importance of the AKT-*like* kinase in the survival of *T. cruzi* and *T. brucei.*

## Results

### Putative allosteric inhibition of *T. cruzi* AKT-*like* protein promotes the generation of apoptosis-like under conditions of nutritional stress

We previously described a putative allosteric inhibitor of the PH domain of the AKT-*like* kinase of trypanosomatids (UBMC4), obtained by bioinformatic strategies (S1 Fig) [24–26]. The morphological and ultra-structural effects of nutritional stress (substitution of the culture medium with PBS for 6 hours), was evaluated in *T. cruzi* epimastigotes previously exposed to sub-lethal concentrations of UBMC4. Nutritional stress alone did not generate effects on morphology and ultra-structure of the parasite, neither morphological markers of apoptosis-like were found, like DNA damage and mitochondrial or cell membrane integrity (Fig 1A and C). However, the parasites subjected to nutritional stress in the presence of UBMC4 showed evident and progressive alterations such as inactivation of the flagellum, reduction of cell size, alteration of the nuclear structure, chromatin condensation in the nuclear periphery, formation of vacuoles and mitochondrial swelling with loss of the integrity of the kinetoplast and other external and internal morphological changes with formation of possible blebs (Fig 1 B, D, E and F).

**Figure 1.**
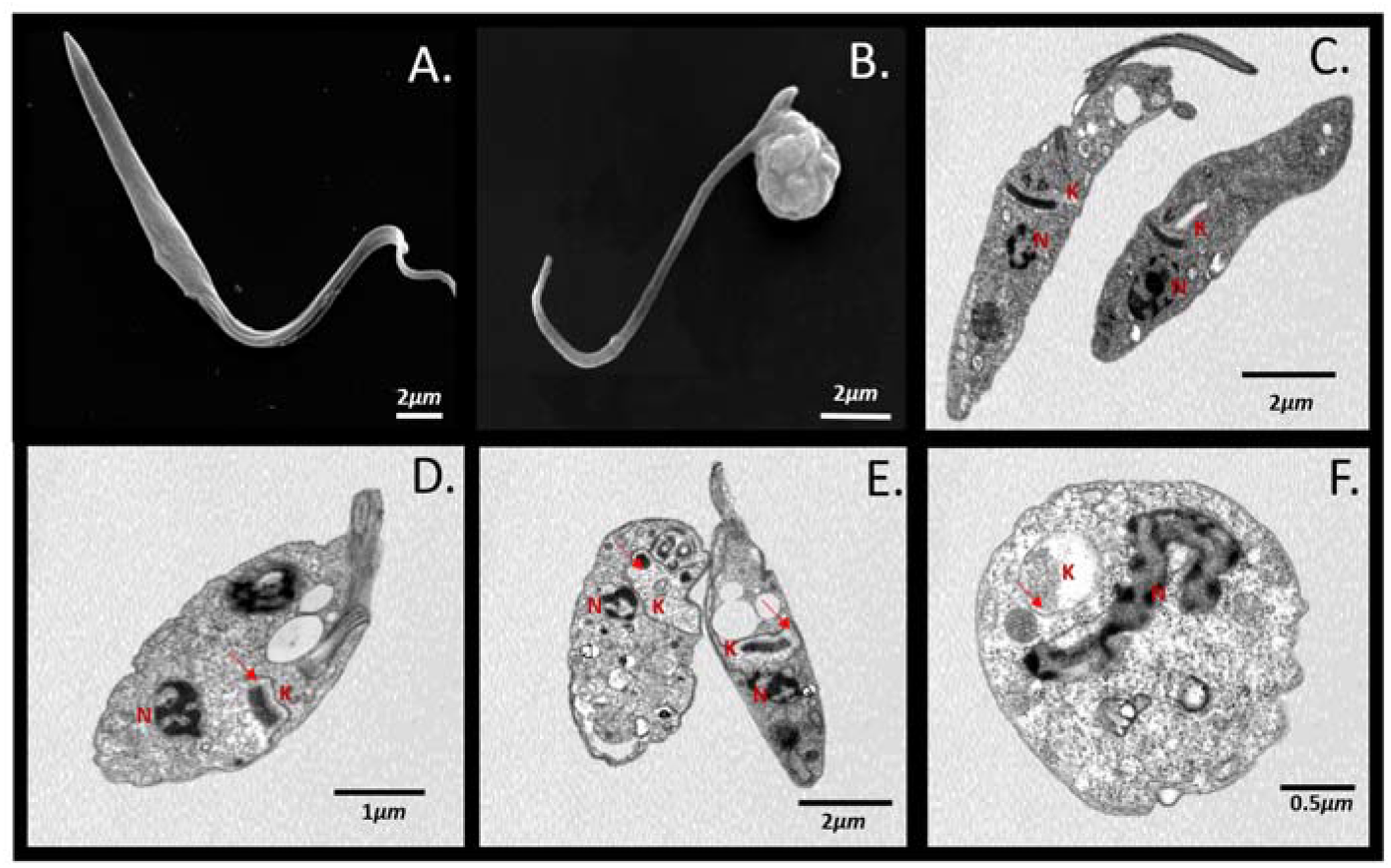
SEM and TEM of *T. cruzi* epimastigotes with nutritional stress and inhibition of their AKT-*like* protein. (A) SEM of epimastigotes with nutritional stress for 6 hours. (B) SEM of epimastigotes with nutritional stress and inhibition of their AKT-*like* protein for 6 hours. (C) TEM of epimastigotes with nutritional stress for 6 hours. (D-E) TEM of epimastigotes with nutritional stress and inhibition of their AKT-*like* protein for 1 hour. (F) TEM of epimastigotes with nutritional stress and inhibition of their AKT-*like* protein for 6 hours. N, Nucleus; K, Cinetoplast; arrows indicate the place of mitochondrial alteration.

Nutritional stress caused parasites to fragment their DNA in 21.9% of the population, (hypodiploids), it promoted mitochondrial depolarization by 22.6% and generated no cell membrane damage in 97.73% of parasite population compared to parasites subjected to normal nutritional conditions (Fig 2 A and B). In turn, the use of 10μM UBMC4 under normal culture conditions did not generate changes in apoptosis-like markers (Fig 2 C). However, its addition in parasite cultures subjected to nutritional stress induced progressive and gradual fragmentation of the DNA up to 39.4% of the population, with an increase of 38.4% with damage to the cell membrane and with 99.9% of the population with depolarization of the mitochondria (Fig 2 D). The effect of the inhibition of AKT-*like* on the mitochondrial polarity combined with nutritional stress was confirmed by performing a kinetic assessment of the mitochondrial potential, showing that nutritionally stressed parasites in presence of the AKT-like inhibitor suffer a mitochondrial depolarization not seen in parasites under normal nutritional conditions (S2 Fig).

**Figure 2.**
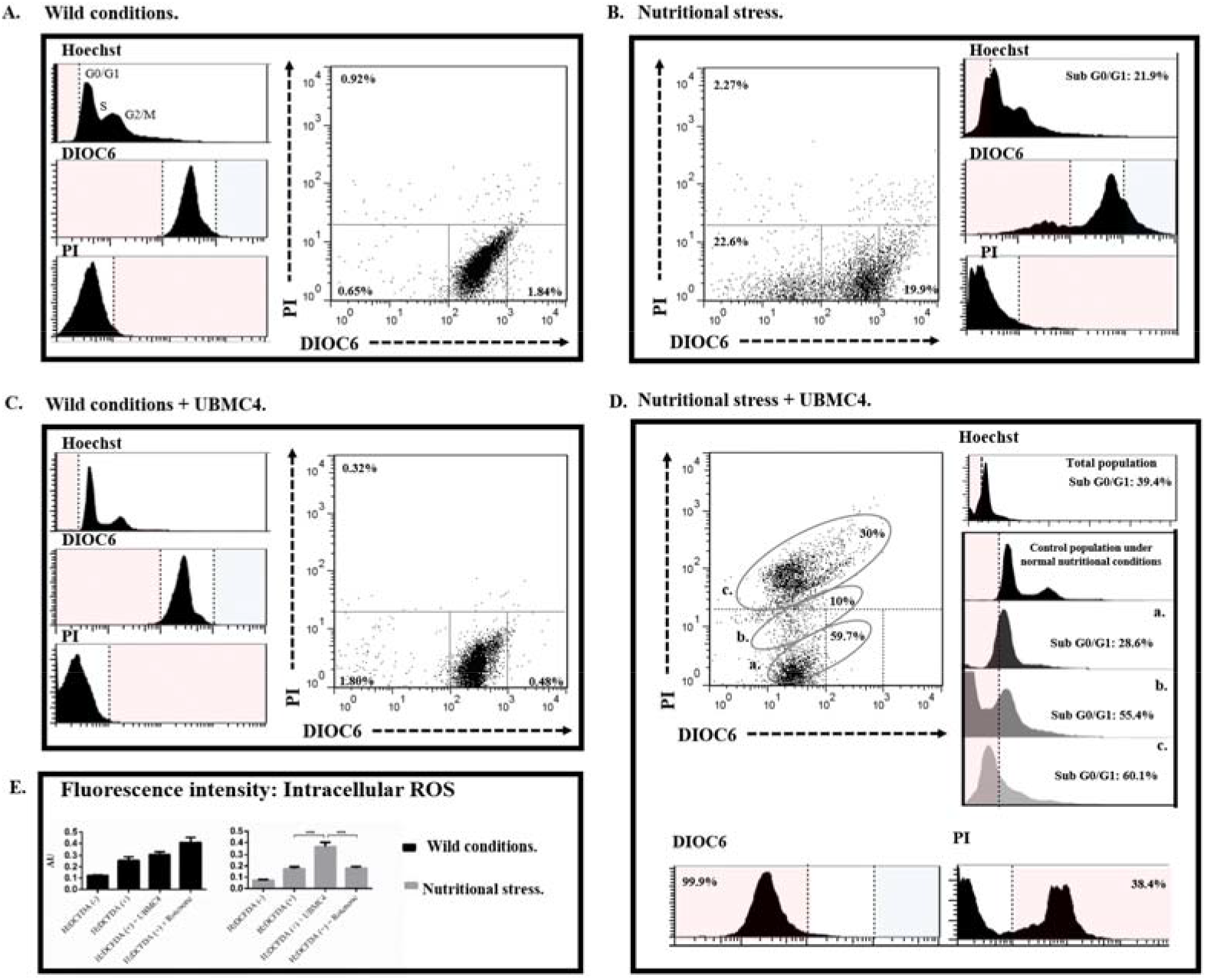
Flow cytometry for the determination of apoptosis-like in *T. cruzi* epimastigotes subjected to nutritional stress and inhibition of their AKT-*like* protein. Frequency histograms for the Hoechst parameters (cell cycle); DIOC6 (ΔΨm); IP (Alteration of the cell membrane) and two-parameter dot-plots for ΔΨm and cell membrane alteration. (A) Epimastigotes under normal nutritional conditions for 24 hours. (B) Epimastigotes under conditions of nutritional stress for 24 hours. (C) Epimastigotes under normal nutritional conditions and inhibition of their AKT-*like* for 24 hours. (D) Epimastigotes under conditions of nutritional stress and inhibition of their AKT-*like* protein for 24 hours. (E) Determination of the intracellular ROS concentration of epimastigotes under normal nutritional conditions and nutritional stress with and without inhibition of their AKT-*like* for 24 hours. The plots represent the trend of three biological replicas per condition and the percentages are equivalent to the average obtained from the three replicas. Asterisks indicate statistically significant differences with respect to control treatments values. (*** p <0.0004).

Similarly, the intracellular ROS concentration of epimastigotes subjected to normal nutritional conditions did not show significant differences between parasites treated and not treated with UBMC4, however, the ROS concentration doubled in parasites treated with the inhibitor and subjected to nutritional stress conditions with respect to the untreated (Fig 2 E).

In the case of *T. cruzi* trypomastigotes, unlike what was observed with epimastigotes, it was found that the inhibition of their AKT-*like* both in normal nutritional conditions and in conditions of nutritional stress promotes increased fragmentation of its DNA, mitochondrial depolarization and the permeabilization of their cell membranes, as well as a progressive change of their morphology. Additionally, the use of 10 μM of chloroquine, a classic autophagy inhibitor, accelerated and increased the effects of UBMC4 with respect to the apoptosis-like markers evaluated (S3 Fig).

### RNAi degradation of *T. brucei* AKT-*like* mRNA promotes apoptosis-like events under conditions of nutritional stress

Gene knockdown by RNAi in the model strain 2T1 of *T. brucei* has been previously reported [27]. We used the same model for the induced degradation of the AKT-*like* mRNA achieving up to 73.13% reduction in the number of transcripts upon induction with tetracycline (*tet*+ parasites) for 48 hours (S4 Fig). AKT-*like* mRNA degradation (*tet*+), did not induce macroscopic changes in the morphology and movement of parasites under normal nutritional conditions (data not shown) compared to transfected parasites without tetracyclin induction (*tet*−), additionally, no significant differences were found in the accumulated growth between parasites with *tet*+ or *tet*− conditions (Fig 3A). On the contrary, *RNAi* induced parasites (*tet*+) subjected to conditions of nutritional stress (elimination of the FBS from the culture medium), presented loss of movement, rounded shape (data not shown) and significant differences in the accumulated growth compared with parasites with no RNAi induction (*tet*−) (Fig 3B). Furthermore, exposure during 48 hours to nutritional stress induced a dramatic growth drop in *tet*+ parasites compared to the *tet*− controls.

**Figure 3.**
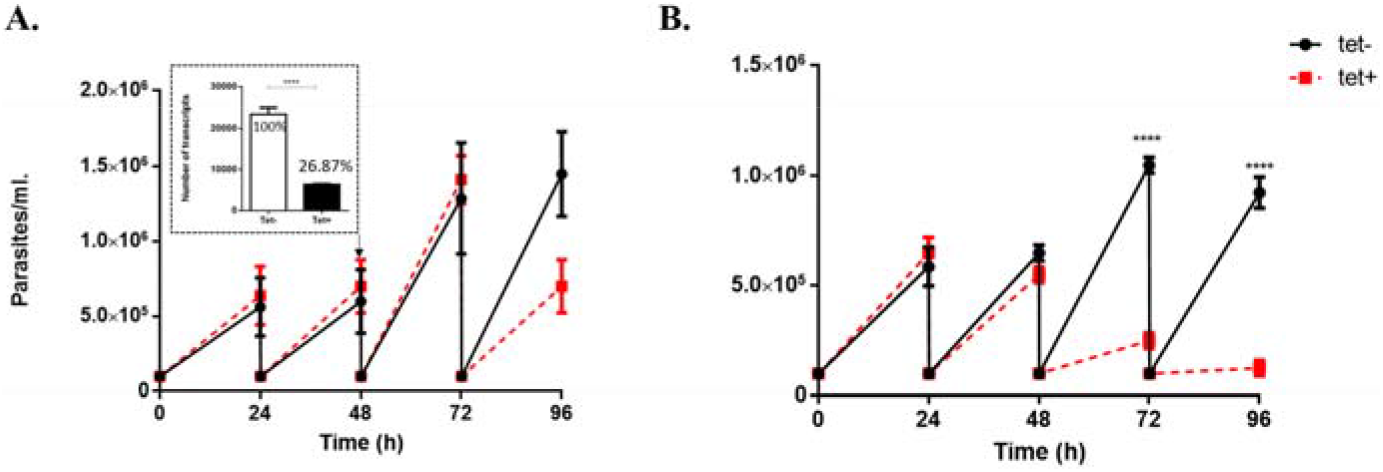
Cumulative growth curves of *T. brucei* blood forms subjected to normal nutritional conditions and nutritional stress under the interference of their AKT-*like* proteins. (A) Growth of parasites with and without RNA interference induction (*tet*+ and *tet*− respectively) under normal nutritional conditions. (B) Growth of parasites with and without RNA interference induction (*tet*+ and *tet*− respectively) under conditions of nutritional stress (elimination of the FBS from the growth medium). The box in Figure A shows the level of overall interference calculated from the relative and absolute quantification of AKT-like transcripts using qRT-PCR (Supplementary Figure 4). Quantifications were made from samples of parasites obtained after 48 hours of induction with tetracycline at 1 μg/ml. The data shown are the results of three independent experimental replicas represented by the means ± their standard deviation, (**** p <0.0001).

Nutritional stress induced progressive DNA fragmentation in *tet*− parasites after 3 hours of treatment, reaching a maximum peak at 6 hours with 42.3% of the hypodiploid population, 74% cells with mitochondrial membrane depolarization and 59% with alterations in their cell membrane with respect to *tet*− parasites cultivated under normal nutritional conditions (Fig 4A, B and E), leaving only 10% of the population without cellular alterations (Fig 4B). *tet*+ parasites under normal nutritional conditions did not generate changes in apoptosis-*like* markers, however, *tet*+ parasites under nutritional stress experienced remarkable apoptosis-*like* events like progressive and gradual DNA fragmentation with an hypodiploid population of 56.3%, mitochondrial depolarization in 84.7% of the population and 54.4% parasites with cell membrane damage (Fig 4C, D and E). Only 5.4% of the population under nutritional stress remain with no alterations in the three apoptosis-*like* markers evaluated.

**Figure 4.**
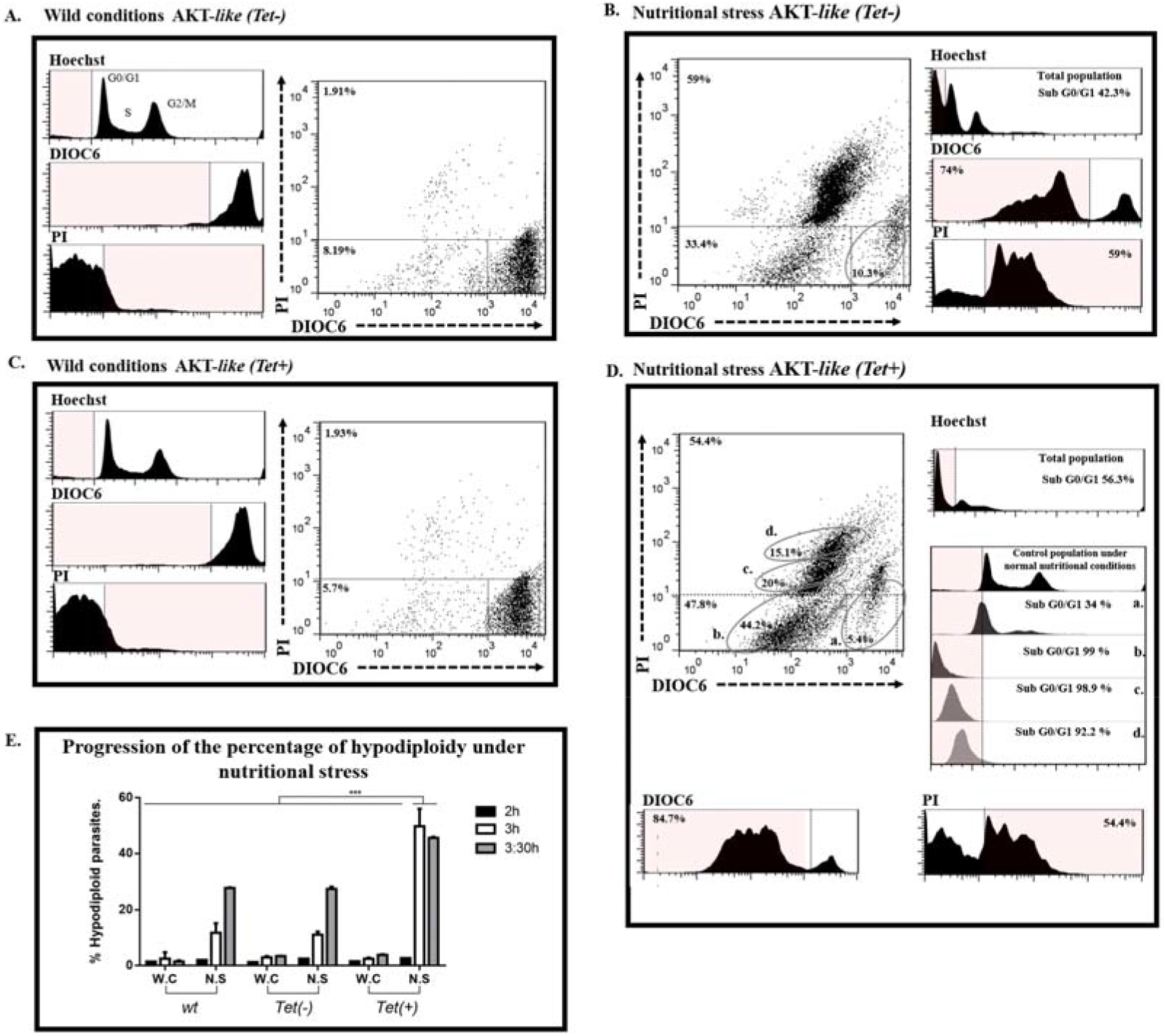
Flow cytometry for the determination of apoptosis-*like* in *T. brucei* blood forms subjected to nutritional stress and low interference of its AKT-*like*. Frequency histograms for the Hoechst parameters (cell cycle); DIOC6 (ΔΨm); IP (Alteration of the cell membrane) and two-parameter dot-plots for ΔΨm and cell membrane alteration. (A) *Tet*− blood forms under normal nutritional conditions for 6 hours. (B) *Tet*− blood forms under conditions of nutritional stress for 6 hours. (C) *Tet*+ blood forms under normal nutritional conditions for 6 hours. (D) *Tet*+ blood forms under conditions of nutritional stress for 6 hours. (E) Percentage progression of hypodiploid parasites subjected to nutritional stress. The interference induction (*tet*+) was carried out 48 hours with 1 μg/ml tetracycline prior to the pressure with nutritional stress. The plots represent the trend of three biological replicas by condition and the percentages progression of hypodiploid parasites are equivalent to the average obtained from the three replicas. Wt; untransfected parasites; tet-, transfected parasites without interference induction; tet+, parasites transfected with interference induction. Asterisks indicate statistically significant differences with respect to the values of control treatments. (*** p <0.0004).

### AKT-*like* knockout in *Leishmania major* promotes apoptosis-like events under conditions of nutritional stress and decreases infectivity of AKT-like −/− parasites

By using the CRISPR-Cas9 system in *Leishmania* parasites [28–30], a double knockout of AKT-*like* in *L. major* was obtained with replacement of both alleles by the neomycin resistance cassette (*AKT-like* −/−) with loss of detectable AKT-*like* transcripts (Fig 5A - B). Generation of double knockout parasites in a single round of transfection indicates that AKT-like gene is not essential for survival of *L. major* promastigotes, at least under controlled nutritional and environmental conditions *in vitro*.

**Figure 5.**
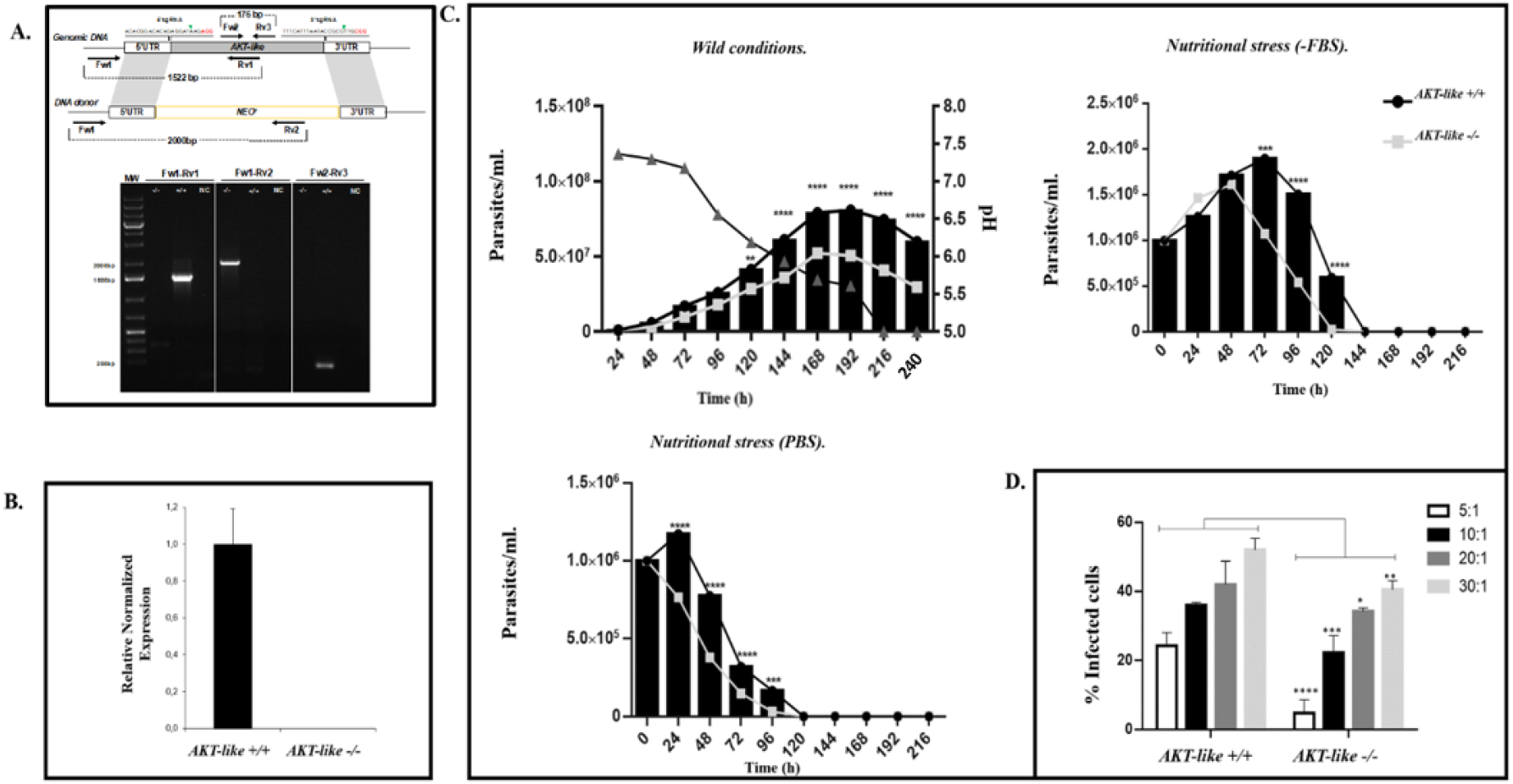
Proof of the AKT-*like* gene knockout and its effect on growth and infectivity of *Leishmania major*. (A) Knockout of AKT-*like*. Up: Schematic representation of the CRISPR/Cas9 strategy used to generate double knockout parasites. AKT-*like* locus (AKT-*like*) was replaced by homologous recombination with a donor DNA cassette containing a neomycin resistance gene (NEO^R^) flanked by 30 nt homology regions from 5’UTR and 3’UTR of the target locus after DNA cleavage by sgRNA-Cas9 complexes. Primers (arrows) used to verify the deletion of AKT-*like* gene and the expected sizes of the PCR-amplified products are indicated. Down: PCR products visualized on agarose gel. An expected 1.5 kb band was obtained using primer pair Fw1-Rv1 in unmodified parasites (+/+), whereas no amplification was observed in knocked-out parasites (−/−). The amplification of the expected 2 kb product using primer pair Fw1-Rv2 only in *AKT*−/− parasites confirmed the replacement with the neomycin resistance gene in both AKT-like alleles. Primer pair Fw2-Rv3 amplify a 176 bp fragment (used for quantification of AKT-*like* transcripts by qRT-PCR) and test for the presence of the AKT-*like* coding sequence. Lane MW shows the DNA marker used. bp, base pairs. NC: Negative control. (B) mRNA was obtained from control (AKT-*like* +/+) and double knockout (AKT-*like* −/−) parasites, and the expression level of the AKT-like gene was analyzed by real-time qPCR. Data represent the means ± SD from 2 independent experiments. (C) Knock-out vs. wild-type parasite growth curves under normal and nutritional stress conditions. (D) Percentage of (*AKT*+/+) and (*AKT*−/−) parasite infectivity. Asterisks indicate statistically significant differences with respect to control treatment values. (**** p <0.0001; *** p 0.0002; ** p 0.0013; * p 0.00365). A value of P <0.05 was considered statistically significant.

No apparent changes were observed in the morphology and movement of the AKT-*like* −/− parasites grown under normal nutritional conditions with respect to wild type parasites *AKT-like* +/+ (data not shown). Additionally, no significant differences were found in the growth curves during the first 96 hours of culture. However, after 120 hours, the *AKT-like* −/− parasites showed significant decrease in growth compared *AKT-like* +/+ cells concomitant with a fall in the culture pH from 7.36 to 5.0 (Fig 5C). It was also observed that the conditions of nutritional stress (both the replacement of the culture medium with PBS and the elimination of the FBS from the culture medium), increased the death rate in AKT*-like* −/− parasites with respect to *AKT-like* +/+ parasites (Fig 5C). This observation was confirmed by conducting cell viability tests under conditions of nutritional stress, showing that AKT*-like* −/− parasites are more sensitive to nutritional stress (S5 Fig). It was also found that AKT*-like* −/− parasites significantly decrease their infective capacity with respect to *AKT-like* +/+ parasites, with an infective concentration 50 (IC_50_) of 9.59 and 5.6 parasites respectively (Fig 5D).

Nutritional stress induced apoptosis-like events after 20 hours in *AKT-like* +/+ parasites evidenced by a progressive and gradual DNA fragmentation with 3.73% of the hypodiploid population, 26.39% cells with mitochondrial depolarization and 22.6% with alterations in their cell membrane integrity as compared with *AKT-like* +/+ parasites under normal nutritional conditions (Fig 6A and B), leaving 70% of the population without apoptosis-like alterations (Fig 6B). On the other hand, AKT*-like* −/− parasites showed no changes in the evaluated markers under normal nutritional conditions, however, when subjected to nutritional stress, those parasite exerted significant differences in the fragmentation of their DNA with respect to *AKT-like* +/+ parasites with an increase in the hypodiploid population of 16.4%, higher percentage of parasites with depolarized mitochondrial membrane (45.3%) and a greater damage in the cell membrane (39%) (Fig 6C, D and F), leaving just 53.1% of the population with no apoptosis-like alterations. We observed an evident difference between *AKT-like* +/+ and AKT*-like* −/− regarding the presentation of apoptosis-like events under conditions of nutritional stress (Fig 6E). The progression of damage over time was confirmed for each parameter at 12 and 24 hours (S6 Fig).

**Figure 6.**
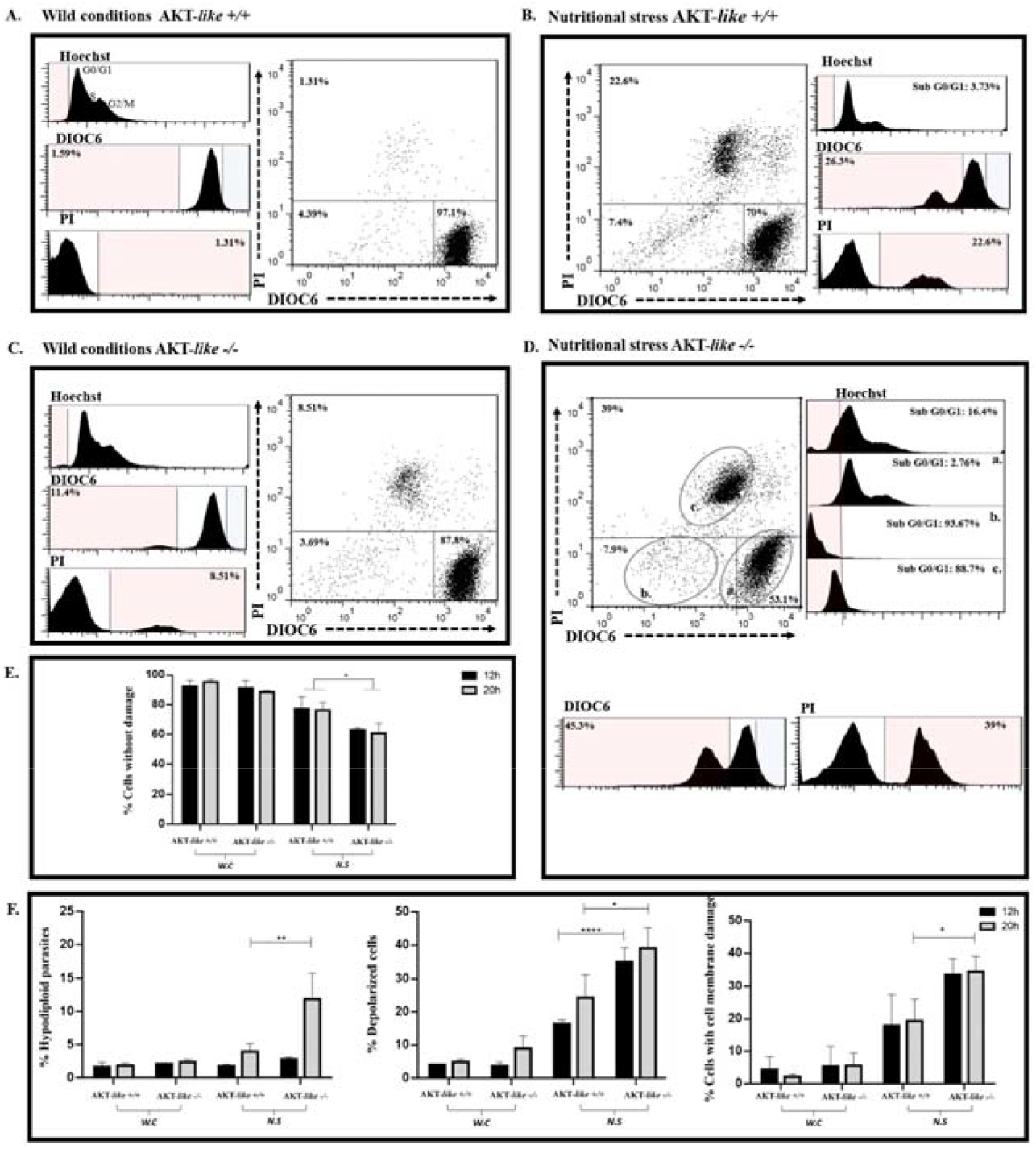
Flow cytometry for the determination of apoptosis-like in *L. major* promastigotes subjected to nutritional stress and the knockout of its AKT-*like*. Frequency histograms for the Hoechst parameters (cell cycle); DIOC6 (ΔΨm); IP (alteration of the cell membrane) and two-parameter dot-plots for ΔΨm and cell membrane alteration. (A) Promastigotes AKT-*like*+/+ under normal nutritional conditions for 20 hours. (B) Promastigotes AKT-*like*+/+ under conditions of nutritional stress for 20 hours. (C) Promastigotes AKT-*like*−/− under normal nutritional conditions for 20 hours. (D) Promastigotes AKT-*like*−/− under conditions of nutritional stress for 20 hours. (E) Percentage of parasites without cell damage (alteration of the apoptosis-*like* parameters selected). W.C; Wild conditions, N.S; Nutritional stress, (* P 0.0111). (F) Percentage of hypodiploid parasites (** P 0.0069); with mitochondrial membrane depolarization (* P 0.0290; **** P <0.0001); and damage to the cell membrane (* P 0.0142). A value of P <0.05 was considered statistically significant.

## Discussion

Tritryps are subjected to different types of stress during their complex life cycle and they must deal with these conditions in order to infect, proliferate and survive. Here, we show by means of three independent experimental approaches that the impairment of the function of the AKT-*like* kinase in Tritryps induces loss of tolerance to nutritional stress, promoting the appearance of apoptosis-like events and reducing their infective capacity. We show that parasites whose AKT-*like* kinase has been inhibited or silenced, suffer DNA fragmentation, mitochondrial damage and loss of cell membrane integrity under stress conditions compared to wild type parasites. Furthermore, we present evidence that deletion of AKT-like gene in *L.major* affects the infective capacity of parasites with respect to unmodified cells where AKT-*like* kinase is present.

*T. cruzi* can affect the host for decades or years with the preexistence of amastigotes inside the tissue. Intercommunication between the parasite and cytosol components of parasitized cells is crucial for the spread and maintenance of the infection, however, little is known about how the parasite manages to survive for long periods without killing its host cell. To achieve this, *T. cruzi* promote the survival of its host cell through strategies such as the production of proteins like the parasite-derived neurotrophic factor (PDNF), a membrane surface trans sialidase which induces the activation of the AKT kinase of the host cell [31]. Nevertheless, the role of the AKT-*like* kinase of the Tritryps, and its participation in resistance to nutritional, immunological and thermal stress events is unknown. Its putative function has been extrapolated from those described in humans, where AKT enzyme is responsible for the activation of cell survival processes by inhibiting apoptosis and maintaining homeostasis [32]. Before the availability of genomic data, the AKT-*like* of *T. cruzi* was first described as an enzyme with kinase activity and putative PKB function. However, its biological function or its importance as a molecular target was not described. Although previously it had been described that AKT-*like* of *T. cruzi* lacked PH domain, unlike human AKT kinases, this hypothesis was later rejected based on computational analysis [33].

The development of a putative allosteric inhibitor (UBMC4) of the *T. cruzi* AKT-*like*, (S1 Fig), allowed us to observe that treated epimastigotes were more sensitive to nutritional stress challenges with respect to untreated parasites, showing a pattern of morphological and ultra-structural alterations typical of apoptosis-like processes, such as cell shrinkage, blebs-like structure formation and damage in the DNA, nuclear structure and mitochondria (Fig 1) [34,35].

The apoptosis markers determined by flow cytometry, revealed that the inhibition of AKT-*like*, promotes greater DNA fragmentation, loss of the integrity of the cell membrane and mitochondrial depolarization in conditions of nutritional stress, where AKT carries out its function as reported for other species (S2 Fig). Membrane depolarization was associated with mitochondrial membrane alterations and increased concentration of intracellular ROS (Fig 2). We interpreted those events as consequences of increased apoptosis-like conditions, revealing that AKT-*like* is crucial for the response to nutritional stress in epimastigotes of *T. cruzi*. It is interesting to note that the trypomastigotes were more sensitive to the putative inhibition of their AKT-*like* protein, even under normal nutritional conditions. In addition, it was observed that the concomitant inhibition of autophagy by chloroquine with the inhibition of AKT-*like*, increases the magnitude of the damage, revealing the interconnection between the autophagy pathway and the PI3K/AKT-*like*/mTOR pathway in trypomastigotes. Our data also suggest that the parasite could activate constitutively the pathways of response to stress conditions where AKT-*like* plays a fundamental role for survival (S3 Fig).

It has been reported that the reduction of about 83% of transcripts of the *S. mansoni* AKT, promotes not only the decrease in the final concentration of the protein, but also significantly decreases the amount of detectable phospho-activated protein, generating defects in glucose metabolism and cell homeostasis [36]. We observed that silencing of AKT-*like* gene in blood forms of *T. brucei* using RNAi resulted in a significant reduction (73.13%) of its transcription level (S4 Fig), which generated defects in the accumulated growth of the parasites when they were subjected to nutritional stress conditions (Fig 3B). However, similar to observed in epimastigotes of *T. cruzi*, the RNAi blood forms of *T. brucei* subjected to normal nutritional conditions, showed no defects in their accumulated growth (Fig 3 A), supporting the hypothesis that the activation and function of AKT-*like* protein is given in response to stress events. In that sense, the apoptosis-like markers showed that the RNAi blood forms of *T. brucei* had fragmentation of their DNA, damage to their cell membrane and mitochondrial depolarization (Fig 4).

Therefore, and despite the fact that the *T. brucei* AKT-*like* had already been interfered in works of massive characterization of kinases by high-performance techniques, the AKT-*like* was never considered as a relevant protein for biological processes such as survival, differentiation, cell cycle and stress response in murine models [37–39]. In our work, we observed that AKT-*like* protein plays an important role in the response to nutritional stress by maintaining DNA integrity, mitochondrial and cell membrane stability.

Previously we probed the presence of an AKT-*like* in various species of the genus *Leishmania*, which is phospho-activated under conditions of thermal and nutritional stress, in addition, it was observed that parasites previously stressed and subsequently challenged with putative inhibitors of human AKT, promoted increased apoptosis-like events and decreased their ability to infect macrophages [24–25]. In accordance with these results, we observed that the knockout of the AKT-*like* of *L. major,* promotes the decrease of its growth rate under normal nutritional conditions from day 5 of culture. This pattern was associated with the increased sensitivity to stress, correlating with the acidification of the culture medium. We determined that AKT*-like* −/− parasites accelerated the decrease of their population when they were subjected to nutritional stress conditions. Interestingly, and in agreement with the results obtained previously by Varela and Tirado 2018, we found that the AKT*-like* −/− parasites were less infective with respect to *AKT-like* +/+ parasites, with an IC_50_ of 9.59 and 5.62 respectively, demonstrating not only the central role that AKT-*like* has in the response to stress conditions, but also in infectivity processes.

We have found a similar pattern of response to nutritional stress in *L. major AKT-like −/−, T. cruzi* and RNAi *T. brucei* parasites evidenced by loss of the integrity of their DNA, mitochondrial depolarization and permeabilization of their cell membrane, suggesting a common function of this protein in the Tritryps (Fig 6 and S6 Fig), where the loss of function leads to increased apoptosis-*like* under nutritional stress conditions.

Finally the data presented here lead us to propose a general model of the function of the AKT-*like* of *T. cruzi*, *T. brucei* and *L. major*, where the main role of the AKT-*like* is the reprograming of cell survival through the inhibition of the apoptosis-like under stress conditions. This protection is achieved in the parasite through the activation of DNA repair pathways, activation of energy metabolism, maintenance of redox balance and inhibition of non-canonic apoptosis-like inducers. Therefore the putative inhibition of the PH domain and the interference or knockout of the AKT-like gene leads to the loss of the kinase function, with the concomitant increase in its vulnerability to nutritional stress and the activation of apoptosis-like death mechanisms. In light of our work we finally defend that the understanding of this process demonstrates the potential of the *AKT-like* as a promising therapeutic target present across the different Tritryp genera, the causing agents of thousands of deaths and disabilities around the world.

## Materials and Methods

### Molecular modeling of *T. cruzi* AKT-*like* protein structure and molecular Docking with UBMC4

In the absence of an experimental three-dimensional structure of the *T. cruzi* AKT-*like*, its amino acid sequence (Q4D6D3 Uniprot) [40], was used as input for the development of a computational predictive model, using the I-TASSER [41] web server processing approach. The kinase model together with the PH domain were subjected to quality tests through the analysis of Ramachandran graphs, as well as the energy curves calculated on the *Swiss Model* server [42]. The model was minimized with gradient descent using the minimization protocol available with UCSF Chimera version 1.10 [43].

The molecular structure of the PH domain’s putative allosteric inhibitor, called UBMC4, was obtained from the ZINC database [24, 44], subsequently, the characterization of its physicochemical properties was performed using filters to determine candidates for drugs proposed in prior works and in compliance with Lipinski’s rules [24, 45]. The *T. cruzi* AKT-*like* model was uploaded to the Ligsite web server to predict a list of potential binding sites with structures of interest [46]. A region surrounding the PH domain was selected and both the protein and the compound were parameterized using AutoDock tools [47]. This process involved the addition of hydrogen bonds to the polar side chains and the estimation of partial charges using the Gasteiger method.

In the Docking strategy, the compound was considered flexible based on the active torsion bonds present in the 3D structure, subsequently, the Docking protocol was executed with *AutoDock Vina* [48], using 20 internal replicas. Interaction scores were predicted in Kcal/mol and molecular interactions were checked graphically to determine the number and type of interaction.

### Cell lines and parasite strains

To determine the *in vitro* cytotoxicity of the UBMC4 compound, human liver cells HepG2 (ATCC code: HB-8065), and human colorectal epithelial carcinoma cells CaCo2 (ATCC code: HTB-37) were cultured, and maintained under the same conditions at 37°C in DMEM medium with 5% FBS and CO2 atmosphere of 5%. In addition, monocyte-derived human macrophages (huMDM) were obtained as described in Daigneault et al., 2010 [49].

For the evaluation of IC_50_ and trypanocidal activity of UBMC4 on *T. cruzi*, the β-galactosidase-producing Tulahuen strain was used, maintained in NNN medium modified with liquid phase (8.5g of NaCl; 10g of glucose; 1L of H20MQ) at 26° C. Additionally, the human promonocytic cell line U937, (ATCC code: CRL1593.2) was cultured in RPMI-1640 medium with 10% FBS, at a concentration of 2.5x 10^5^ cells/ ml with 5% CO_2_ atmosphere with an incubation temperature of 37° C [50].

Epimastigotes of the *T. cruzi* Gal 61 strain were kept at 28° C in RPMI 1640 medium supplemented with 0.02% hemin, 0.5% peptone trypticase, 0.002M HEPES and 10% inactivated FBS and a mixture of antibiotics (100 U/mL−1 penicillin and 0.1 mg/mL−1 streptomycin). The parasites were subcultured weekly in exponential growth phase [51,52].

African green monkey kidney epithelial cells, VERO (ATCC code: CCL-81), were grown in DMEM culture medium with 10% FBS at 37°C and with a 5% CO_2_ atmosphere and maintained until a confluence of 90%, at which time the 10% FBS DMEM medium was replaced by DMEM FBS-free medium, which allowed to obtain a growthless monolayer for infection tests with metacyclic trypomastigotes of a Gal61 *T. cruzi* strain and obtaining blood stage trypomastigotes [53]. Epimastigotes grown to the stationary phase were used to perform these infections at a ratio of 3:1 parasites per cell, with a total concentration of 2×10^6^ cells. Metacyclic trypomastigotes and cells were maintained at 37°C with 5% CO_2_ atmosphere in DMEM medium without FBS. After a period of 10 to 12 days, the blood stage trypomastigotes that emerged were harvested [53].

The blood forms of *T. brucei* 2T1 strain, which constitutively expresses the element repressor protein (TerR) and has a landing platform for exogenous expression constructs in the intergenic region that codes for the tubulin of chromosome 2a [54], were grown at 37°C with a 5% CO_2_ atmosphere in HMI-9 culture medium supplemented with 10% inactivated FBS [27].

Promastigotes of *Leishmania major* Friedlin (AKT-*like* +/+) and its genetically modified double Knockout derivative (AKT-*like* −/−) were grown at 26°C in Schneider medium, supplemented with 20% FBS.

### RNA interference line (RNAi) for *Trypanosoma brucei* AKT-*like*

An RNAi cell line of *T. brucei* AKT-*like* was obtained, using the tetracycline-inducible system (Tet) of the 2T1 strain developed by Alsford et al. in 2005 [54,55–59]. PRpa^isL1/2^ plasmid was used in dsiRNA generation for knockdown of *T. brucei* AKT-*like* protein (Tb427.06.2250) [27].

For the construction of the plasmid, genomic DNA of 1×10^8^ blood stage was extracted for conventional PCR amplification with a Q5^®^ High-Fidelity DNA polymerase (New England Biolabs) from the interference inserts corresponding to the sense and anti-sense fragments of a portion of 429 bp AKT-*like* genomic sequence of *T. brucei*. These inserts were amplified by using oligonucleotides designed with the TrypanoFAN bioinformatics tool: https://dag.compbio.dundee.ac.uk/RNAit/. The amplification conditions were those recommended by the manufacturer and after obtaining the digested and purified inserts and plasmids, parallel and independent ligation reactions were performed. Subsequently, the respective plasmids cloned with the individual sense and anti-sense inserts were taken and a second round of digestions and ligations (sub cloning) were performed under the same conditions to clone the opposite inserts and obtain the double-cloned final constructs. The transfection process was conducted as suggested by Alsford, et al., [54,55–59] and cloning was verified by sequencing.

The gene knockdown was carried out by inducing 1×10^4^ parasites/ml with tetracycline at a final concentration of 1 μg/ml. All subsequent experiments were performed 48 hours post induction. To verify the differential expression of the AKT-like transcripts in the interfered parasites, the qPCR technique was carried out using the Verse 1-step RT-qPCR SYBR Green ROX kit (Thermo Scientific) and the protocols recommended by the manufacturer of the LightCycler^®^ 96 termal cycler (Roche). Differential transcription was determined by both, absolute and relative quantification methods as reported in the LightCycler^®^ 96 System user manuals. For the absolute quantification method, a standard curve was constructed with the pJET1.2/blunt vector cloned with a fragment of the 1336 bp AKT-like sequence and using the 18S House Keeping gene as normalizer. To determine the effect of knockdown on the growth of parasites, accumulated growth curves were performed [60,61].

### Double *knockout* cell line for the AKT-*like* of *Leishmania major*

To obtain the double-knockout cell line for AKT-*like* (LmjF.30.0800), *L. major* parasites were edited using CRISPR-Cas9 system, described previously by Beneke, et al, 2017. Primers were designed using online resource LeishGEdit (http://www.leishgedit.net). The donor DNA was amplified from pTNeo plasmid using the primer pair forward 5’- CTCGACAGCCGGAAAGCACAGTAGGCACTCGTATAATGCAGACCTGCTGC-3’ and reverse 5’-TGATAGAGAGAGAGAGGCGTGATCAGCTCGCCAATTTGAGAGACCTGTGC-3’, to generate a PCR product containing a neomycin resistance gene flanked by 30 nt homology regions from 5’UTR and 3’UTR of AKT-like locus. Single-guide RNA (sgRNA) templates, including a T7 promotor for *in vivo* transcription, were amplified using common primer G00 (sgRNA scaffold) and specific primers for 5′sgRNA (5’-GAAATTAATACGACTCACTATAGGACACGGACACAGAGGATAAGGTTTTAGAGCTAGAAATAGC-3’) and 3’sgRNA (5’GAAATTAATACGACTCACTATAGGTTTCATTTAATACCGCGTTGGTTTTAGAGCTAGAAATAGC-3’), which lead to DNA cleavage close to start and stop codons of AKT-*like* coding sequence, respectively. Log-phase *L. major* promastigotes expressing Cas9 and T7 RNA polymerase proteins [30] were co-transfected with donor DNA and sgRNA templates using V-033 program of Amaxa Nucleofector System (Lonza). Transfected parasites were selected with 50 μg/ml neomycin and cloned in semi-solid culture medium. Genomic DNA were obtained from several clones, and the replacement of AKT-like gene and correct integration of the neomycin-resistance cassette was evaluated by conventional PCR amplification. Furthermore, the differential expression of AKT transcripts was determined by qPCR-like, both in unedited parasites (AKT-*like*+/+) and in knockout parasites (AKT-*like*−/−) [62].

Growth curves were performed using Neubabuer chamber counts every 24 hours for AKT-*like* +/+ and AKT-*like* −/− parasites under normal and stress growth conditions (FBS or FBS replaced by PBS). Additionally, the culture medium pH was determined every 24 hours and the data obtained were analyzed using the two-way analysis of variance (ANOVA) statistical test for multiple comparisons with a significance of 95% through the GraphPad Prism 8 program. Evaluation of cell viability was performed for the AKT-*like* +/+ and AKT-*like* −/− cell lines through the fluorometric technique by staining with Resazurin at 0.15 μg/ml [63].

*In vitro* infectivity (percentage of infection and infective capacity) of the AKT-*like* +/+ and AKT-*like* −/− lines on U937 pro-monocytes (CRL-1593-2TM) differentiated to macrophages, was conducted using the methodology described in [64]. The effect of the initial concentration of parasites versus parasite load was determined by analyzing data in the GraphPad Prism 8 program using non-linear regression and two-way analysis of variance with a significance of 95%.

### *In vitro* studies of the trypanocidal and cytotoxic activity of UBMC4

The U937 cell line was differentiated to macrophages by incubating them with phorbol 12-myristate 13-acetate (PMA) at 0.1 μg/ml. After this, the cells were adhered to 96-well plates for 72 hours by adding 100 μl of cells to the final concentration of 0.25×10^6^ cells/ml in each well. Subsequently, stationary phase epimastigotes of the β*-galactosidase*-producer Tulahuen strain were added in a proportion of 5 parasites/cell and incubated 24h at 37°C with 5% CO_2_ [50]. To calculate the IC_50_, 6 serial dilutions of UBMC4 were made starting with an initial concentration of 200 μM, and benznidazole (BZ) at 20 μg/ml was used as a control for the experiment’s effectiveness. Additionally, a parasite viability control (untreated cells) was included in the experiment.

The cells in the process of infection were incubated for 72 hours, after which the medium was removed and 100 μl of the β*-galactosidase* substrate (chlorophenol red β-d-galactopyranoside CPRG) at 100 mM and 0.1% Nonidet P-40 were added to each well incubating at 37°C for 3 hours. The colorimetric change was read using Thermo Scientific’s ELISA BariosKan Flash dish reader, at an excitation length of 570nm. The percentage of inhibition or IC_50_ was calculated by the Probit J.D method using the GraphPad Prism 8 statistical program.

Cytotoxicity and selectivity index were evaluated according to the ability of UBMC4 to cause the death of the huMDM, obtained as described in Tirado et al., 2018 [24] while in the HepG2 and CaCo2 cell lines the cytotoxicity was evaluated, by the MTT method (3-(4,5-dimethylthiazol-2-yl) 2,5-diphenyltetrazole) described elsewhere [44,65,66].

### *In vitro* evaluation of UBMC4 inhibitor activity on *T. cruzi* epimastigotes in the presence or absence of nutritional stress conditions

Both mild (FBS deprivation from the RPMI medium) and strong nutritional stress (total deprivation of nutrients by replacing the RPMI medium with PBS) were selected to determine the effects of putative inhibition of *T. cruzi* AKT-*like*, for which the qualitative and quantitative determination of the effects of stress and treatment with UBMC4 on the generation of apoptosis-like events was performed. To evaluate apoptosis-like events, markers such as morphological changes in the structure and ultra-structure determined by scanning and transmission electron microscopy, alterations of mitochondrial membrane potential, changes in the intracellular concentration of ROS, as well as DNA and cell membrane damage, determined by flow cytometry, were carried out.

### Scanning and transmission electron microscopy

Epimastigotes of *T. cruzi* treated and untreated with UBMC4, under normal and stressed nutritional conditions, were fixed for two hours with 2.5% glutaraldehyde in 0.1M cacodylate buffer. The parasites were adhered for 10 minutes on coverslips previously covered with 0.1% poly-L-lysin and subsequently washed with cacodylate buffer and sequentially dehydrated with 30%, 50%, 70% and 90% acetone (v/v) during 5 minutes on each incubation. After this, the cells were critically dried with CO_2_ and attached to the heel of the scanning electron microscope (SEM). Finally, the samples were coated with a 20nm thick gold layer and observed in a Jeol JSM6010Plus-LA scanning electron microscope at 20kv. In the case of transmission electron microscopy, the parasites were fixed for two hours with glutaraldehyde and cacodylate buffer, after which the samples were treated for one hour with 1% OsO4 diluted in the same buffer. The parasites were dehydrated as before and then embedded in PolyBed-812 resin. Ultrathin sections were collected in copper grids and stained with 5% aqueous uranyl acetate and lead citrate. After this, the samples were examined in a Jeol JEM-1400plus transmission electron microscope, operating at 80kv.

### Analysis of the intracellular production of ROS in epimastigotes of *T. cruzi* treated with UBMC4

The intracellular ROS concentration was determined by fluorometric tests with the Diacetate 2’, 7’-dichlorodihydrofluorescein (H2DCFDA) probe at 0.2 μg/ml, for which 1.4×10^6^ epimastigotes under normal nutritional and stressed nutritional conditions were treated with 10μM of UBMC4 for 24 hours. As controls, both treatments without the probe and cells that were labeled only with H2DCFDA were performed to determine baseline fluorescence. As a control, the parasites were subjected to the challenge with 50mg/ml of rotenone (a ROS inducer) for one hour prior to labeling and reading.

After the treatments, the parasites were washed and resuspended in 100 μl of the solution with H2DCFDA and transferred to 96-well plates protected from light and incubated for one hour at 37°C. The test was read in an Elisa plate reader at an excitation wavelength of 495nm and emission of 527nm. Each treatment was performed in triplicate and the data were reported in arbitrary units of fluorescence [67].

### Analysis of apoptosis-*like* in *T. cruzi*, *T. brucei* and *Leishmania major* by flow cytometry

The analysis of apoptosis-*like* was carried out by triple staining with 3,3’-dihexyloxacarbocyanine (DIOC6); propyl iodide (PI) and Hoechst from Thermo Fisher Scientific, to determine the change in mitochondrial membrane potential (ΔΨ*m*); damage to the cell membrane and the cell cycle respectively [24,68]. All readings were performed on a BD FACSCanto II 4/2/2 Sys IVD flow cytometer and the data obtained were analyzed using the FlowJo7.6.2 software. The differences between treatments were determined by two-way analysis of variance (ANOVA), which allowed us to make multiple comparisons with a significance of 95% using the GraphPad Prism 8 statistical program. In all cases, 20 μl of parasites (1×10^7^/2ml) were stained per treatment with 300 μl of Hanks balanced saline and 10 μl of the staining mix, containing 10 μl of IP, 0.5 μl of DIOC6 and 0.5 μl of Hoechst, at a concentration of 5 μg/ml, 70 nM, and 10 mg/ml respectively. Subsequently, the parasites were incubated at room temperature for 15 minutes until reading.

Epimastigotes of *T. cruzi* were grown in normal nutritional conditions, and after this, the exponential phase parasite culture was used to perform nutritional stress treatments, as well as the challenge with both the UBMC4 inhibitor and the control drug Bz. The experiments were performed in 24-well plates at an initial concentration of 1×10^7^ total parasites in 2ml/well. Nutritional stress was induced eliminating the RPMI medium and replacing it with PBS. The treatments evaluated were parasites in complete culture medium, and in PBS (with and without 10μM of UBMC4) for 24 hours at 28°C in three independent biological replicas. The 24 well plate, was centrifuged at 3000 rpm for 5 minutes and washed once with PBS buffer, after which parasites were resuspended in a final volume of 2ml for marking and reading on the cytometer.

In addition to the apoptosis markers, an action kinetics test of UBMC4 was performed using the DIOC6 probe, for which epimastigotes in complete culture medium and nutritionally stressed for 24 hours, were read continuously for a period of 18.3 minutes. After the first minute of reading, DIOC6 (1μl at 5μg/ml per treatment) was added and read continuously until the membrane potential curve became stabilized, which took approximately 7 minutes. After the stabilization of the base curve that represents the basal potential of the mitochondrial membrane, a challenge with 20 μM of UBMC4 was performed 12 minutes after the start of the reading. The alteration of the curve’s slope was interpreted as: positive, hyperpolarization; and negative, depolarization.

For the *T. cruzi* trypomastigotes, the same markers mentioned above were used. A total of 1×10^6^ trypomastigotes were centrifuged at 2000 rpm for 5 minutes, eliminating the culture medium. After this, they were reconstituted in a DMEM medium with 10% FBS or only PBS as a condition of nutritional stress. The treatments evaluated were: parasites cultured in normal and nutritional stress conditions challenged with and without UBMC4 at 10μM; Chloroquine at 10μM and UBMC4/Chloroquine at the same concentrations for two- and five-hour periods. All experiments were contrasted with Bz as a control compound.

The same markers were evaluated in the *T. brucei* blood forms of the 2T1 strain in exponential phase with and without induction of their AKT-*like* interference (48 hours before induction with tetracycline at 1 μg/ml) under normal and nutritional stress conditions for a period of 6 hours with three independent biological replicas. In this case, the experiments were carried out in 24-well plates in 2ml at a concentration of 1×10^5^ parasites/ml. Blood forms were then counted, centrifuged at 3000 rpm/10 minutes and washed with PBS before being reconstituted with the final volume at the mentioned concentration of parasites with the desired treatment. In this case, the nutritional stress selected was the elimination of the FBS since in previous qualitative analyzes it was determined that these parasites are highly susceptible to the elimination of all the nutrients from the medium, making it impossible to carry out the experiments with strong nutritional stress.

For the determination of apoptosis-*like* in *Leishmania major*, 5×10^7^ exponential phase parasites of AKT-*like* +/+ and AKT-*like* −/− clones were subjected to normal and nutritional stress conditions for periods of 12, 20 and 24 hours after which they were dyed and read as mentioned for *T. cruzi* and *T. brucei*.

### Statistical analysis

The experimental development of each section was carried out through the application of three biological replicas and three experimental replicas for each data obtained. In all cases, analysis of variance or ANOVA was applied for multiple comparisons after determining that the data were normal, or in the opposite case, if the data showed non-parametric distribution, the Kruskal-Wallis statistical test was applied.

## Author contributions

**Conceptualization**: Rubén E. Varela-M, Marcel Marín-Villa

**Data curation:** Andres Felipe Diez Mejia, Rubén E. Varela-M, Marcel Marin-Villa,

**Formal analysis:** Andres Felipe Diez Mejia, Rubén E. Varela-M, Marcel Marin-Villa,

**Funding acquisition:** Rubén E. Varela-M, Marcel Marín-Villa

**Investigation:** Andrés Felipe Díez Mejía, Maria Magdalena Pedroza, Lina M. Orrego.

**Methodology:** Mauricio Rojas’ Lina M. Orrego, Maria Clara Echeverry, Sergio Pulido, Maurilio José Soares, Sara M. Robledo, Marcel Marín-Villa, Rubén E. Varela-Miranda, Carlos Enrique Muskus, José María Pérez-Victoria.

**Project administration**: Marcel Marín-Villa

**Resources:** Maria Clara Echeverry, Sara M. Robledo, Maurilio José Soares, José María Pérez-Victoria.

**Supervision:** Marcel Marín-Villa

**Visualization:** Rubén E. Varela-M, Andrés Felipe Díez Mejía

**Writing – original draft:** Andres Felipe Diez Mejia, Rubén E. Varela-M, Marcel Marin-Villa.

**Writing – review & editing:** Andres Felipe Diez Mejia, Lina M. Orrego, Rubén E. Varela-M, Marcel Marin-Villa, Sergio Pulido, Sara M. Robledo, Carlos Enrique Muskus.

## Acknowledgments

Rodrigo Ochoa for bioinformatic analysis.

## Supporting Information

**S1 Fig.**
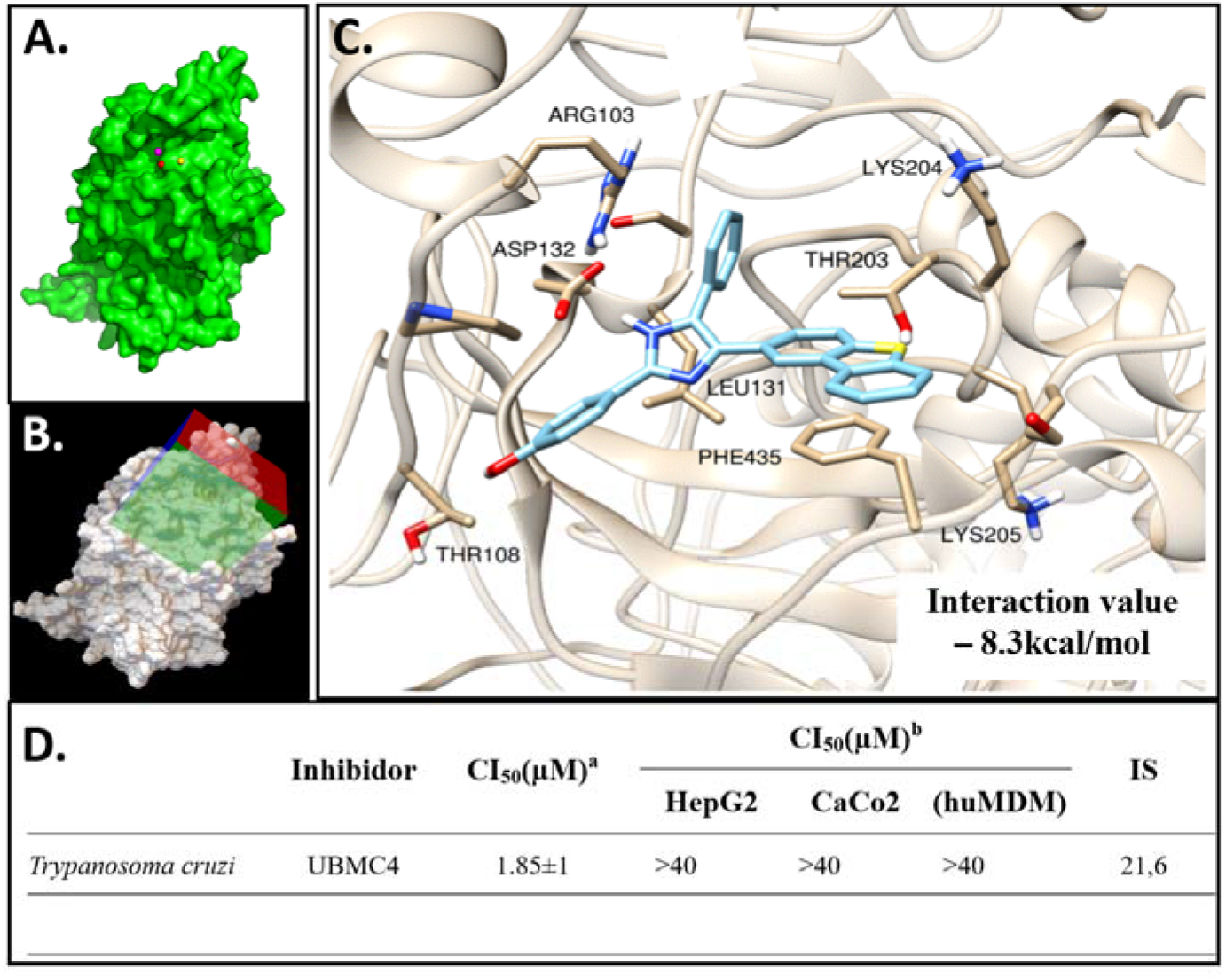
(Related to Figures 1 and 2). Structural modeling and molecular dynamics of the AKT-*like* protein in *T. cruzi*. (A) Structural model of the AKT-*like* protein in surface format. The pocket predicted by MetaPocket 2.0 server tools are presented with colored spheres. (B) Search box configured in the AutoDock Tools program for molecular docking of the predicted pocket near the PH domain of the protein with ~640.000 compounds in 3D format, executed on the DrugDiscovery@TACC server (drugdiscovery.tacc.utexas.edu). (C) Pocket interacting with UBMC4. The key residues that undergo some kind of molecular interaction with the compound and the interaction energy obtained are shown. (D) Cytotoxic and trypanocidal activity of the UBMC4 compound. ^**a**^ IC_50_, inhibitory concentration of 50 for T. cruzi amastigotes. ^**b**^ IC_50_, lethal concentration of 50 for macrophage derived monocytes (huMDM); HepG2 line liver tissue cells and colon tissue line (CaCo2). ^**c**^ IS, selectivity index between (huMDMD) and T. cruzi amastigotes. The data shown represent the average value of the experiments ± the standard deviation.

**S2 Fig.**
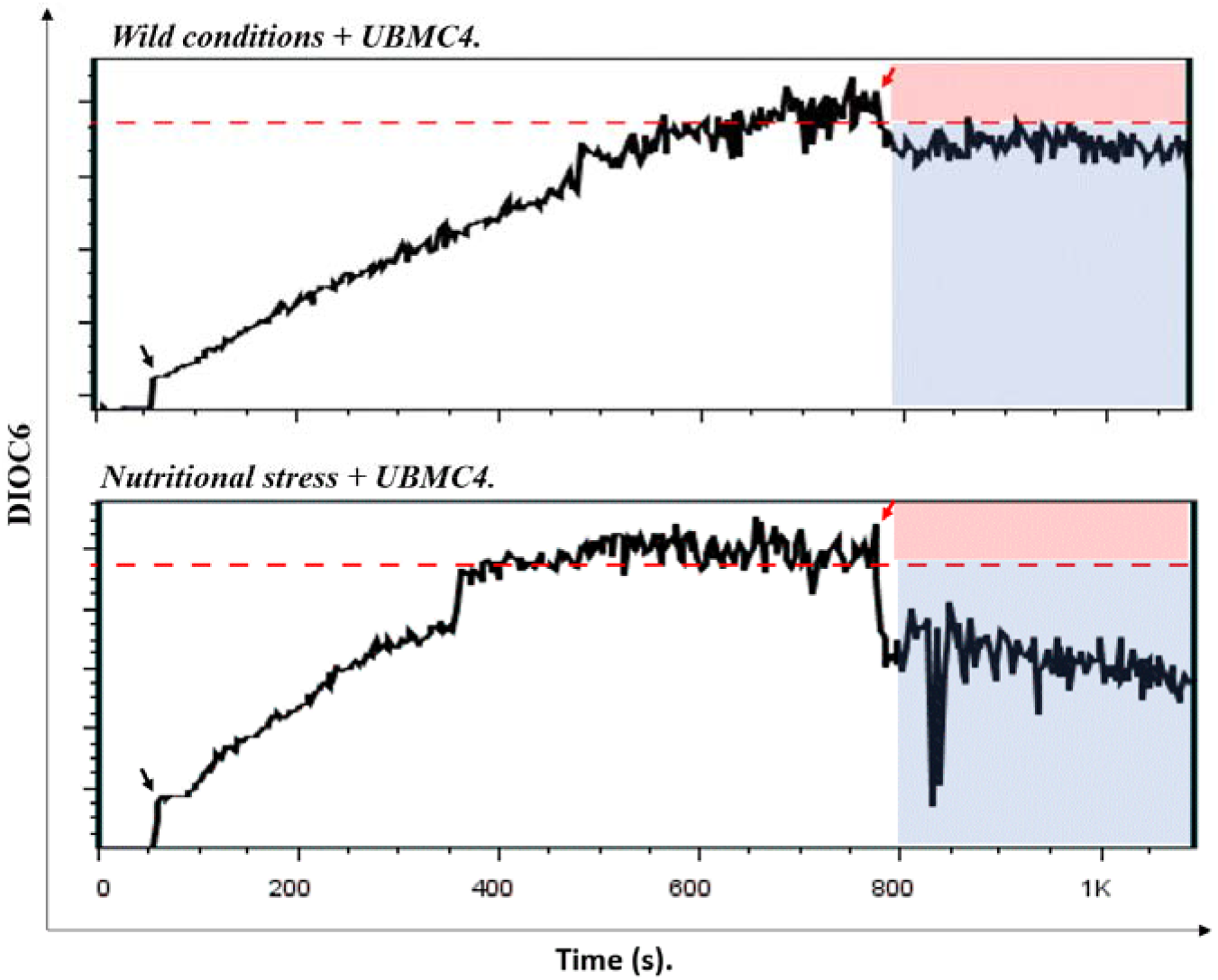
(Related to Figure 2). Kinetics of mitochondrial membrane potential of *T. cruzi* epimastigotes subjected to normal and nutritional stress conditions under inhibition of their AKT-*like* protein. The upper box shows the Ψm kinetics of *T. cruzi* epimastigotes under normal nutritional conditions and the inhibition of their AKT-*like* protein. The bottom box shows the kinetics of parasites subjected to nutritional stress conditions and the inhibition of their AKT-*like*. The kinetics was evaluated for a period of 18.3 minutes. The red boxes show the area interpreted as mitochondrial hyperpolarization and the blue ones, the area interpreted as mitochondrial depolarization.

**S3 Fig.**
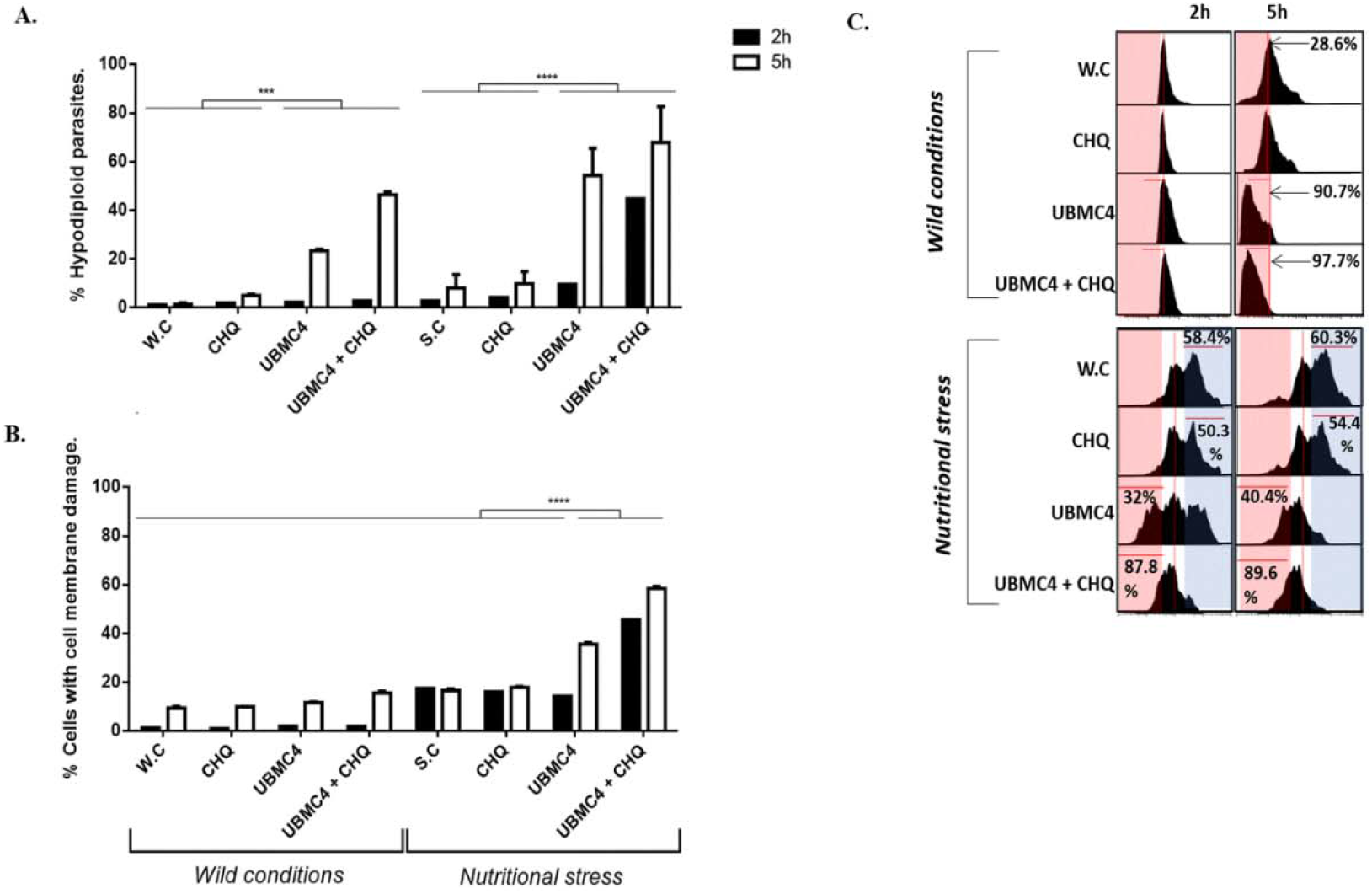
(Related to Figures 1 and 2). Determination of apoptosis-*like* in *T. cruzi* trypomastigotes subjected to nutritional stress and inhibition of their AKT-*like*. The effects of AKT-*like* inhibition and autophagy were determined by flow cytometry and TEM by the use of chloroquine in *T. cruzi* trypomastigotes. (A) Percentage of *T. cruzi* trypomastigote hypodiploidy under normal nutritional conditions and nutritional stress for 2 and 5 hours. (B) Percentage of damage in the cell membrane of *T. cruzi* trypomastigotes under normal nutritional conditions and nutritional stress for 2 and 5 hours. (C) Frequency histogram of the Ψm of *T. cruzi* trypomastigotes under normal nutritional conditions and nutritional stress for 2 and 5 hours. W.C, wild conditions; CHQ, Chloroquine; S.C, conditions of nutritional stress. Asterisks indicate statistically significant differences with respect to the values of control treatments. (**** p <0.0001).

**S4 Fig.**
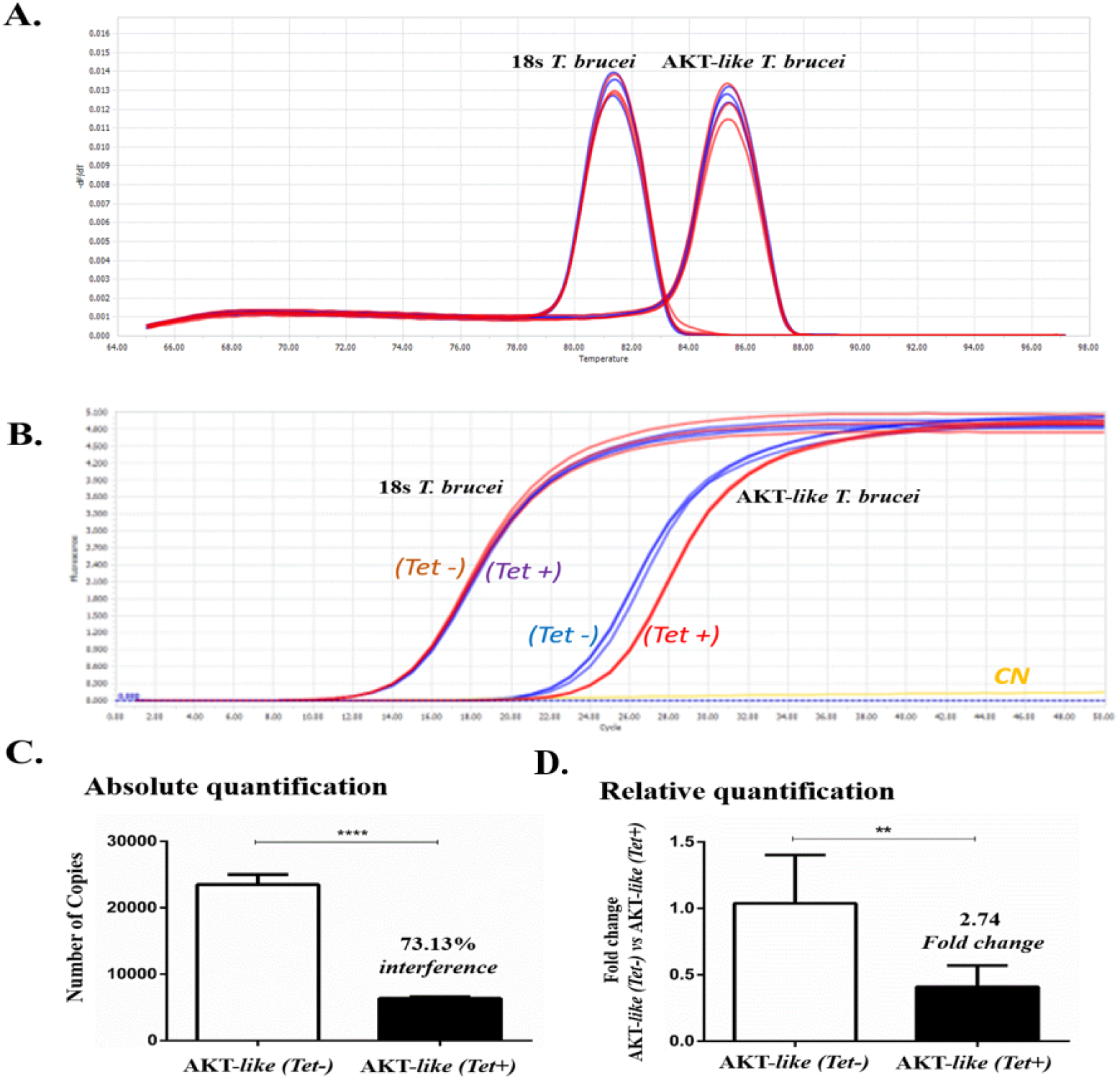
(Related to Figures 3 and 4). Quantification by qRT-PCR of the interference level in AKT-*like* transcripts of *T. brucei*. (A) Melting curves for verification of amplification specificity. (B) Amplification curves from the transcripts of the 18S and AKT-*like* genes of parasites with and without interference induction for 48 hours. (C) Quantification of number of copies or transcripts of the AKT-like determined by absolute quantification analysis of parasites with and without interference induction for 48 hours. (D) Quantification of the rate of change in the AKT-like transcripts determined by relative quantification analysis of parasites with and without interference induction for 48 hours. The data represents the average ± the standard deviation of 3 independent experimental replicas. Asterisks indicate statistically significant differences with respect to the values of control treatments. (**** p <0.0001; ** p <0.0012).

**S5 Fig.**
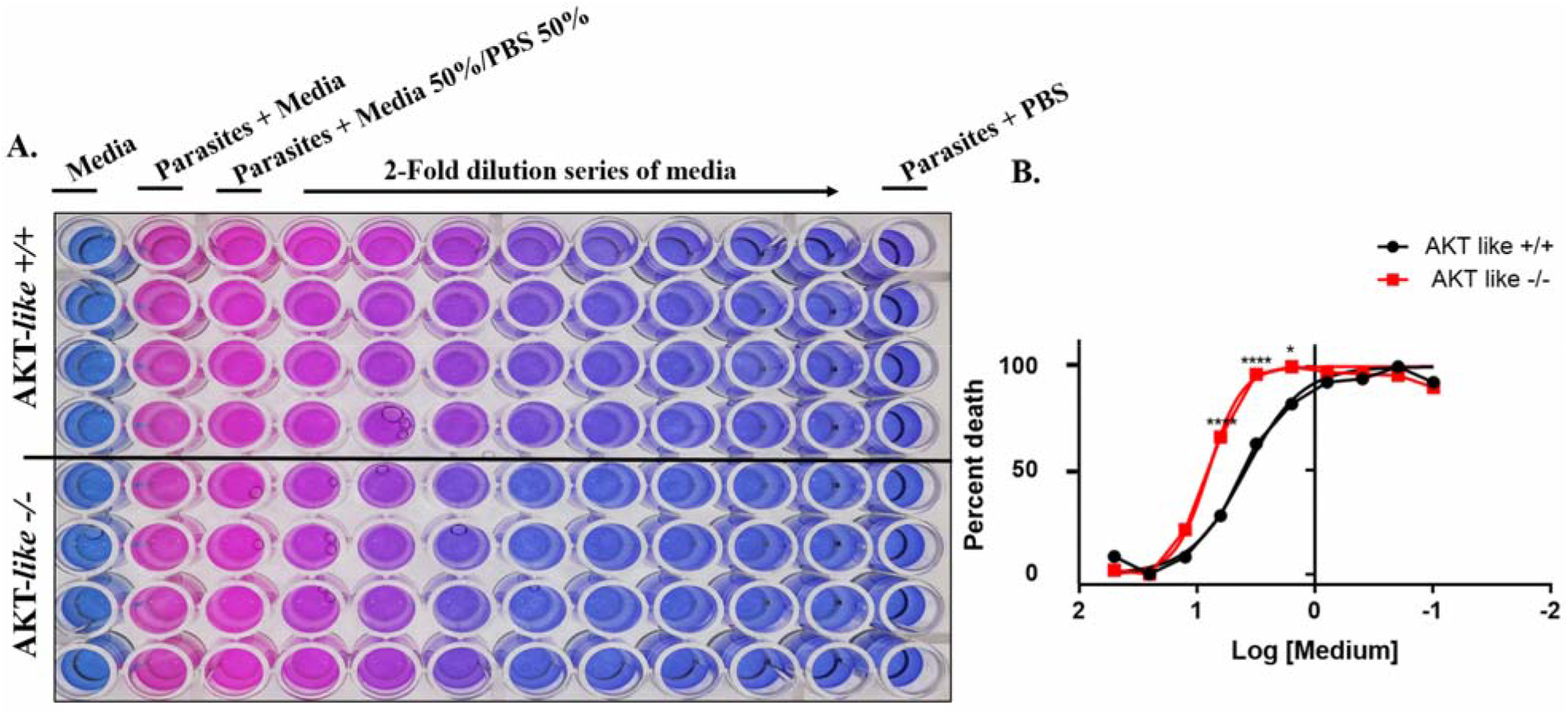
(Related to Figures 5 and 6). Cell viability of *L. major* promastigotes subjected to nutritional stress and the knockout of their AKT-*like*. Cell viability test by fluorometry of Resazurin metabolism. (A) AKT-*like* +/+ and AKT-*like* −/− promastigotes subjected to conditions of nutritional stress by cultivation in descending concentrations of complete environment. (B) Dose response analysis (Average percentage/Percentage of cell death), (* P 0.0362; **** P <0.0001).

**S6 Fig.**
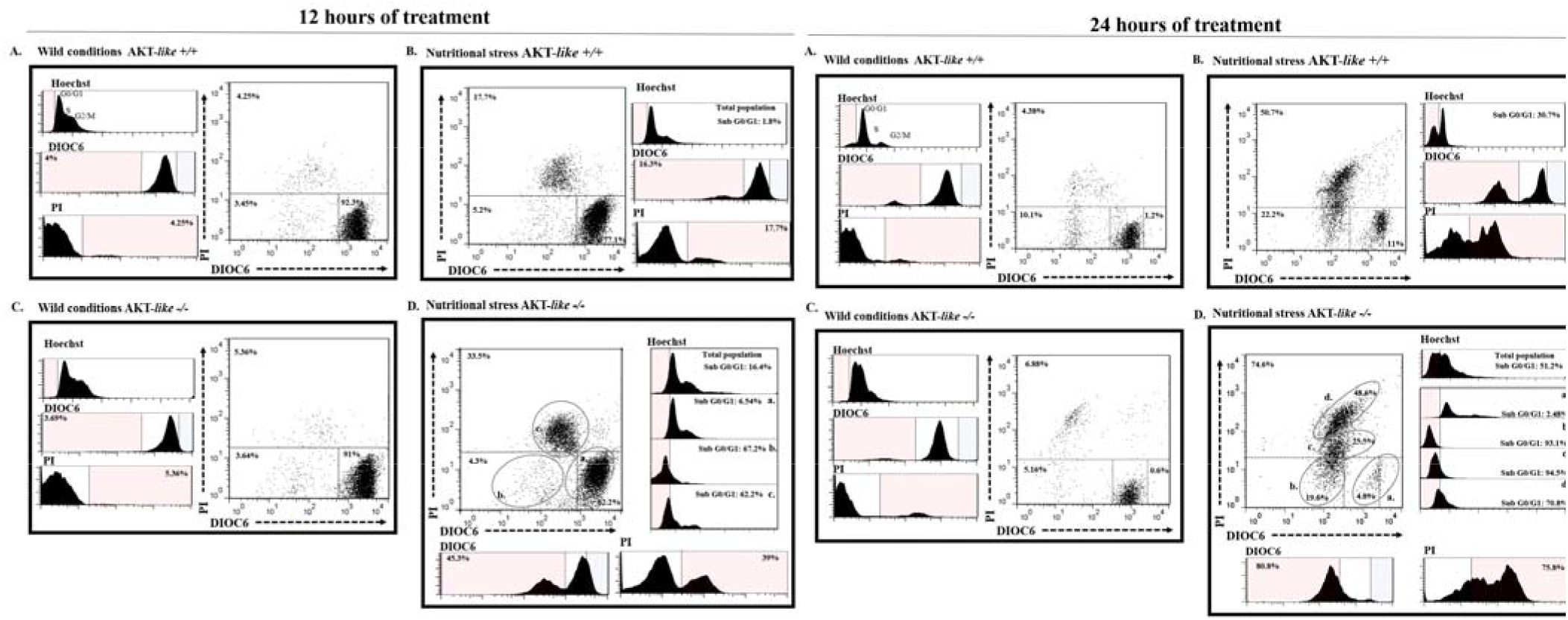
(Related to Figures 5 and 6). Flow cytometry for the determination of apoptosis-*like* in promastigotes of *L. major* subjected to nutritional stress and the knockout of its AKT-*like* in 12- and 24-hour periods. Frequency histograms for Hoechst parameters (Cell cycle); DIOC6 (ΔΨm); IP (Alteration of the cell membrane) and two-parameter Dot-plots for ΔΨm and alteration of the cell membrane. (A) AKT-*like* +/+ promastigotes under normal nutritional conditions for 20 hours. (B) AKT-*like* +/+ promastigotes under conditions of nutritional stress for 20 hours. (C) AKT-*like* −/− promastigotes under normal nutritional conditions for 20 hours. (D) AKT-*like* −/− promastigotes under conditions of nutritional stress for 20 hours.

